# Ultrafast Power Doppler Ultrasound Enables Longitudinal Tracking of Vascular Changes that Correlate with Immune Response After Radiotherapy

**DOI:** 10.1101/2023.08.04.552076

**Authors:** Shannon E. Martello, Jixin Xia, Jiro Kusunose, Benjamin C. Hacker, McKenzie A. Mayeaux, Erica J. Lin, Adrienne Hawkes, Aparna Singh, Charles F. Caskey, Marjan Rafat

## Abstract

**Background:** While immunotherapy shows great promise in patients with triple negative breast cancer, many will not respond to treatment. Radiotherapy has the potential to prime the tumor-immune microenvironment for immunotherapy. However, predicting response is difficult due to tumor heterogeneity across patients, which necessitates personalized medicine strategies that incorporate tumor tracking into the therapeutic approach. Here, we investigated the use of ultrasound (US) imaging of the tumor vasculature to monitor the tumor response to treatment.

**Methods:** We utilized ultrafast power doppler US to track the vascular response to radiotherapy over time. We used 4T1 (metastatic) and 67NR (non-metastatic) breast cancer models to determine if US measurements corroborate conventional immunostaining analysis of the tumor vasculature. To evaluate the effects of radiation, tumor volume and vascular index were calculated using US, and the correlation between vascular changes and immune cell infiltration was determined.

**Results:** US tumor measurements and the quantified vascular response to radiation were confirmed with caliper measurements and immunostaining, respectively, demonstrating a proof-of-principle method for non-invasive vascular monitoring. Additionally, we found significant infiltration of CD8^+^ T cells into irradiated tumors 10 days after radiation, which followed a sustained decline in vascular index and an increase in splenic CD8^+^ T cells that was first observed 1 day post-radiation.

**Conclusions:** Our findings reveal that ultrafast power doppler US can evaluate changes in tumor vasculature that are indicative of shifts in the tumor-immune microenvironment. This work may lead to improved patient outcomes through observing and predicting response to therapy.

## Introduction

The incidence of breast cancer is rising, with over 300,000 new diagnoses annually [1]. Approximately 15% of cases are identified as triple negative breast cancer (TNBC), which is characterized by a disproportionally high death rate due to its aggressive nature and lack of targeted therapies [2,3]. Radiotherapy (RT) is a mainstay of TNBC treatment and significantly reduces the risk of recurrence after surgery [4–6]. RT exerts biological effects beyond tumor cell killing on both short-and long-term scales, altering physical and molecular vascular characteristics and influencing immune cell infiltration and phenotype [7–10]. Recently, it has been suggested that RT can be used as a tool to elicit immune cell infiltration and activation in the tumor, thereby improving the efficacy of immunotherapies in otherwise immunologically “cold” tumors [11–13]. While some TNBC patients respond well to immunotherapies like immune-checkpoint blockade [14–16], it remains difficult to predict therapeutic outcomes due to varying degrees of malignancy, immune landscape, and tumor heterogeneity across patients.

The tumor vasculature influences the therapeutic response molecularly and physically by controlling immune cell trafficking, anti-tumor immunity, and drug delivery. Increases in tumor microvessel density over time and compared to healthy tissue are indicative of disease progression [17–20], providing a metric to monitor response to therapy. For example, a reduction in vessel density is known to occur within days to weeks after RT exposure [21,22], and a rebound in vascular growth has been shown to occur prior to a detectable change in tumor volume in recurrent tumors [23]. Recent work has also linked vascular changes in skin cancer with immune cell infiltrates and response to immunotherapy, demonstrating a critical connection between tumor vasculature and the immune system [24]. Non-invasive monitoring of the tumor vasculature post-RT therefore has potential not only to determine disease progression and treatment response but also to predict outcomes of immunotherapy.

Previous studies have used contrast-enhanced ultrasound (US) to differentiate vessel sizes and measure fibrosarcoma response to radiation [23,25] and high frequency power doppler US to monitor breast tumor response to RT and anti-angiogenic therapy [26]. Here, we use ultrafast power doppler US to measure tumor vasculature and track changes over time. Ultrafast US combines improved resolution and sensitivity, allowing for deeper penetration of tissues and more accurate reconstruction of vascular changes over time [27,28]. In this study, ultrafast US images were taken at multiple timepoints following tumor irradiation in two TNBC models with varying metastatic capacity to investigate vascular dynamics and immune cell infiltration after RT. We found that vascular changes following RT can be captured by US imaging and that the vascular indices measured by US correlate with vessel density quantified by traditional methods at the same timepoint post-RT. We also found that a decrease in US-derived vascular index post-RT preceded an increase in CD8^+^:CD4^+^ T cell ratios, demonstrating the potential of this non-invasive vascular imaging tool for examining therapy-induced shifts in the immune microenvironment.

## Methods

### In vivo mouse models and radiation

Animal studies were performed in accordance with the institutional guidelines and protocols approved by the Vanderbilt University Institutional Animal Care and Use Committee. Female BALB/c mice (8-10 weeks old) were inoculated with 5x10^4^ 4T1 or 1x10^6^ 67NR murine TNBC cells into the right #4 mammary fat pad. The number of tumor cells inoculated in each model was determined based on previous studies [29,30], which allowed for robust tumor growth while allowing us to carry out experiments without tumor ulceration at endpoint. Tumor length and width were measured using digital calipers (Fisher Scientific) beginning one (4T1) or two (67NR) weeks post-inoculation. Tumor volume was estimated by volume = (L_1_ ^2^xL )/2, where L_1_ is the smallest and L_2_ is the largest measurement of the tumor. At 100 mm^3^, tumors were irradiated to 12 Gy using a 300 kVp cabinet x-ray system filtered with 1.57 mm aluminum. During irradiation, mice were anesthetized with 2.5% isoflurane and shielded using a 2 cm thick Cerrobend shield with 1 cm wide and 1.5 cm long apertures to expose the tumors. Transmission through the shield was less than 1%.

### US imaging

Animals were anesthetized with 1.5-2% isoflurane. An imaging array (L12-5, Philips, Reedsville, PA) was acoustically coupled to the palpable tumor target through a 6 cm tall cavity filled with warm degassed water. A Verasonics Vantage Ultrasound system (Verasonics Inc, Kirkland, WA, USA) was used for imaging planning and acquisition. Real-time B-mode and power doppler imaging were used to check coupling quality, locate tumor center, and identify the spatial extent of the tumor in the elevational direction. Microbubbles were made in-house using established techniques and administered via retro-orbital injection [31]. To capture a volumetric image of the tumor, 6-12 image planes were acquired with 1 mm spacing to capture. For each plane, 5100 frames of radio frequency (RF) data were acquired at frame rate of 510 Hz at 8.9 MHz. Each frame was comprised of 3 angled compounding planes ranging from -18° to 18°. The acquired RF data was reconstructed offline to in-phase and quadrature (IQ) data.

### US offline processing

Motion correction was performed every 30 frames. The mean of each frame was taken and normalized by the 30-frame median. The standard deviation of 15 frames with low mean amplitude was calculated, and a stability threshold was defined as three folds of standard deviation plus the normalized mean. Outlier frames that exceeded the threshold were replaced with linear interpolation from the remaining frames. The motion corrected IQ data was then passed through a singular value decomposition (SVD) filter with 5% cutoff, a Butterworth bandpass filter from 25 Hz to 125 Hz, and a Top-hat filter with 2 mm disk size to help remove static tissue signal and enhance moving signals that depicts vasculature. The filtered data was then summed to produce a power doppler image. The summed power doppler images were passed through Frangi filter that enhanced tubular structures [32]. The tumor region was manually segmented from power doppler images overlayed on the corresponding B-mode images using MATLAB ginput function (The MathWorks, Inc., Natick, Massachusetts, US). Total tumor volume was evaluated by summing the volume for each acquired plane. The dimmest vessel region on the Frangi filtered image was selected manually with MATLAB ginput and a brightness threshold was calculated by taking the mean of the selected region. A binary mask was generated using the brightness threshold, segmenting the vasculature within the tumor. The total vasculature volume was calculated from the number of voxels in the masks across the planes. The vascular index was determined by calculating the ratio of the total vasculature volume to the total tumor volume.

### Immunofluorescence (IF) and immunohistochemistry (IHC)

At 1 or 10 days post-RT, tumors and spleens were resected and placed in 10% neutral buffered formalin (NBF) for 24h at 4°C and then in 70% ethanol until embedding in paraffin. Sections (4 µm) were deparaffinized, rehydrated, and boiled in citric acid (10 mM pH 6) for 30 min for antigen retrieval. For immunofluorescence (IF) detection, blocking in 10% donkey serum was followed by overnight incubation at 4°C with anti-CD31 (1:100; Goat host, R&D Systems AF3628) and subsequent incubation with a donkey anti-goat AlexFluor594 secondary antibody (1:500; Invitrogen A-11058). An autofluorescence quenching solution (Vector Laboratories SP-8400-15) was used before nuclear staining with 4’,6-diamidino-2-phenylindole dihydrochloride (DAPI) and mounted with a coverslip. For chromogenic (IHC) detection, sections were treated with 3% hydrogen peroxide following antigen retrieval and blocking was performed with 10% goat serum, followed by overnight incubation with CD4 or CD8 primary antibodies (1:100; Rat host, Invitrogen 14-0808-82 (CD8), Invitrogen MA1-146 (CD4)). Sections were then sequentially incubated with anti-rat biotinylated secondary antibodies (1:200; goat host, Vector Laboratories BA-9400-1.5), substrate from a DAB peroxidase substrate kit (Vector Laboratories SK4105), and counterstained with hematoxylin 7211 (Epredia Series). A no primary control was performed for all conditions in both fluorescence and chromogenic detection to confirm antibody specificity.

### Immunostaining analysis

All samples were imaged using an inverted Leica microscope (DMi8) with a 20x or 40x objective and a Leica DFC9000GT sCMOS fluorescence or MC190 HD digital camera. A lumencor mercury-free SOLA light engine was used as the illumination source for fluorescence microscopy. DAPI, Texas Red, and Green Fluorescent Protein filter cubes were used to capture nuclei, CD31^+^, and ⍺-smooth muscle actin (⍺SMA)^+^ staining, respectively. For each tissue, a single plane of the entire section was captured using the autofocus and TileScan functions in the Leica LASX imaging software. Fiji was used to analyze all images [33]. For tumor vessel analysis, the area of positive CD31 staining for each tumor type was first identified based on images of the corresponding no primary control sample. Quantification of vessel count per field and the average vessel cross-sectional area per field was conducted in Fiji after each image was thresholded to analyze only areas of positive staining. The areas of positive staining (i.e. vessels) were then counted and their cross-sectional areas were measured using the Fiji Analyze Particles function. For each sample, 10 analyzed images were randomly selected and the vessel count and average vessel cross sectional area per field was plotted. For pericyte coverage analysis, a mask of CD31^+^ staining was created and the fraction of ⍺SMA^+^ staining within the mask was measured to determine the amount of pericyte coverage per field of view. For each sample, 10 analyzed images were randomly selected and the percent of ⍺SMA^+^ staining that made up the CD31^+^ staining mask was plotted. To quantify the number of CD4^+^ and CD8^+^ cells per field in tumor samples, the Fiji cell counter plugin was used. The CD8^+^:CD4^+^ ratio was calculated by averaging the number of CD8^+^ and the number of CD4^+^ cells per field for each mouse and then calculating the ratio. To measure the percent area of CD4^+^ or CD8^+^ staining in spleen samples, the area of positive staining was first identified based on corresponding images of the no primary control. Images were then thresholded and the area fraction of positive staining was quantified in Fiji.

### Flow cytometry

At 1 and 10 days post-RT, tumors, spleens, and lungs were resected and minced in serum-free RPMI media. Tumors and lungs were then placed in a 400 µg/mL DNase I and 20 µg/mL Liberase digestion solution and incubated for 30 minutes in a 37°C water bath. Digestion was stopped by adding RPMI supplemented with 10% fetal bovine serum. Lungs and spleens were then incubated in ACK lysis buffer for 2 minutes before adding 3% fetal bovine serum in phosphate-buffered saline (PBS) to halt lysis. All cells were stained with Aqua fixable viability stain (ThermoFisher) and FC receptors were blocked with CD16/32 (Biolegend) while simultaneously staining for all surface markers. Cells were incubated for 20 minutes at 4°C, rinsed with PBS, and fixed with 1% NBF for at least 20 minutes at 4°C. Flow cytometry was performed on a four-laser Amnis CellStream machine (Luminex) and FlowJo software (Becton Dickinson) was used for analysis. The following antibodies were used for analysis: CD45-BV650 [30-F11] (1:200; rat host, BioLegend 103151), CD11b-APC [M1/70] (1:200; Rat host, eBioscience 17-0112-82), F4/80-PE [BM8] (1:100; rat host, BioLegend 123110), Ly6G-efluor450 [1A8] (1:100; rat host, eBioscience 48966882), GR-1-APC-efluor780 [RB6-8C5] (1:200; rat host, eBioscience 47-5931-82), MHC2-AF488 [M5/114.15.2] (1:160; rat host, BioLegend 107616), CD11c-BV570 [N418] (1:200; Armenian hamster host, BioLegend 117331), CD3-APC [17A2] (1:100; rat host, BioLegend 100236), CD4-AF488 [GK1.5] (1:200; rat host, BioLegend 100423), and CD8-PE [53-6.7] (1:200; rat host, BioLegend 100708).

### Statistical analysis

To determine statistical significance, differences in CD31, CD8, and CD4 expression and in immune cell populations after radiation and between the two tumor types or between timepoints were determined by two-way ANOVA. Paired t-tests were used to determine statistical significance of vascular index between timepoints for each condition. Pearson and Spearman correlation coefficients were used to determine correlation between tumor volumes derived from ultrasound versus caliper measurements. All statistical analyses were performed in GraphPad Prism 10.

## Results

### US tracks differences in tumor volume and vascular index over time

To evaluate whether ultrafast US can non-invasively monitor tumor response to RT, metastatic 4T1 and non-metastatic 67NR tumors were irradiated to 12 Gy—a non-curative dose that allowed for the study of both vascular and immune cell changes over time. At 1 day prior to and 1 and 8 days after RT, tumor volume was quantified using B-mode US, and total vascular volume was quantified using high frame rate doppler US to determine the feasibility of detecting a response on both short and long timescales (**Figure 1A, B**). US-derived tumor volumes were overall highly correlated with manual caliper measurements at each timepoint for 67NR (0 Gy) and 4T1 (0, 12 Gy) (**Figure 1C**). Furthermore, linear regression analysis of each radiation condition demonstrated overlapping 95% confidence intervals of slopes and intercepts (**Supplemental Table S1**), highlighting the ability of US to track with caliper measurements regardless of tumor type and treatment. While the volumes were weakly correlated in irradiated 67NR tumors, US measurements captured the marked stall in 67NR tumor growth after radiation that matched caliper measurements (**Figure 1D**), which may be critical to patient care and prediction of therapy response. The differences in calculated volumes in the irradiated 67NR tumors are likely a result of the large reduction in tumor volume 8d post-RT that led to errors in caliper measurements due to difficulties palpating the tumor, further highlighting the need for an improved method like US to track tumor response during treatment. In contrast to the change in growth of 67NR tumors post-RT, the highly metastatic 4T1 tumors were unaffected by RT. Accuracy of the US volume measurements compared to caliper measurements and the ability to detect major changes in tumor volume post-RT reveals the potential of this technique to provide a more quantitative method for tracking growth changes over time post-RT.

**Figure 1.**
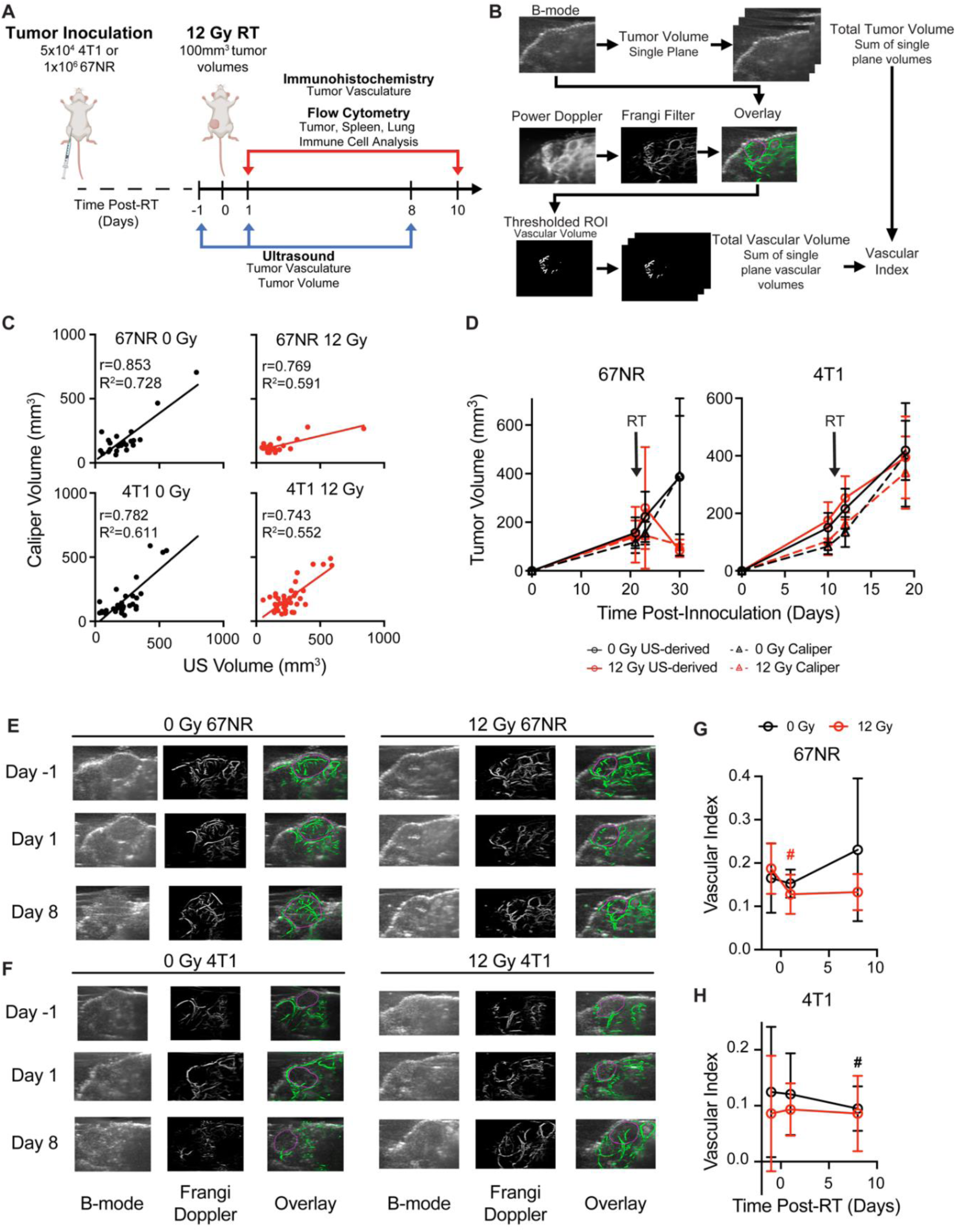
Ultrasound (US) imaging captures vascular changes and accurately tracks changes in tumor volume post-RT. **A.** Experimental timeline. Mice were inoculated with murine 67NR or 4T1 TNBC cells, and tumors were irradiated to 12 Gy once they reached 100mm^3^. US imaging was performed 1 day prior to and 1 and 8 days after RT. Tumors were resected at 1 and 10 days post-RT for immunostaining and flow cytometric analysis. **B.** Overview of imaging and processing to derive tumor volume and vascular index from US measurements. **C.** Correlation of US-and caliper-derived tumor volumes at -1, 1, and 8 days post-RT in 67NR and 4T1 tumors. Pearson correlation (r) and linear regression analyses (R^2^) were performed to determine the association between tumor volumes measured by US and caliper measurements. **D.** Tumor volumes measured via US at -1, 1, and 8 days post-RT (solid lines, circles) compared to manual caliper measurements taken at the same time points (dashed lines, triangles) for 67NR and 4T1 tumors. Error bars show standard deviation. Representative images of B-mode, Frangi-filtered power doppler US, and an overlay of the vascular masks on the tumor volume in **(E)** 67NR and **(F)** 4T1 tumors. Vessels imaged with doppler are shown in green. Purple circles indicate the identified tumor volumes. Quantification of the vascular volume from doppler images and tumor volume from B-mode images enabled calculation of vascular index post-RT over time in **(G)** 67NR and **(H)** 4T1 tumors. Error bars represent standard deviation with *#* denoting p<0.05 in comparison to day -1 timepoint within each dose (black=0 Gy, red=12 Gy) as determined by paired t-tests between timepoints. For each analysis, the number of biological replicates (n) were as follows: 67NR (0 Gy: n=9 for -1/1 day, n=4 for 8 day; 12 Gy: n=8 for -1/1 day, n=4 for 8 day), 4T1 (0 Gy: n=13 for -1/1 day, n=6 for 8 day; 12 Gy: n=15 for -1/1 day, n=9 for 8 day).

Noticeable differences in vascular index between the tumor types and after RT were also observed in US analysis. The vascular index of each tumor was determined by measuring the total vascular volume in Frangi filtered doppler US images and normalizing to the measured tumor volume calculated from B-mode images (**Figure 1B**), enabling non-invasive quantification of vascular changes that are independent of tumor size. US images showed that the 67NR tumors were highly vascular with clear vascular structures visible throughout the tumor volume (**Figure 1E**). Conversely, 4T1 tumors appeared largely avascular throughout the center, and visualization of vascular networks was difficult (**Figure 1F**), indicating a necrotic core. At 1 day post-RT, 67NR tumors showed a significant decrease in vascular index from 0.17 ± 0.04 to 0.13 ± 0.04 that did not recover up to 1 week later (**Figure 1G**). Control mice bearing 67NR tumors showed a 35% increase from 0.17 ± 0.08 to 0.23 ± 0.17 at 8 days post-RT. There were no significant changes in vascular index in the 4T1 tumors in both irradiated and control mice (**Figure 1H**), and 4T1 tumors exhibited a lower vascular index compared to 67NR at all time points.

### Image-derived vascular index correlates with immunostaining analysis

IF staining for CD31 was performed on control and irradiated 67NR and 4T1 tumors to validate and compare the US findings to standard practices of quantifying vascular changes in both pre-clinical and clinical studies [18,34]. 67NR tumors were characterized by higher vessel density compared to 4T1 tumors in both control and irradiated conditions (**Figure 2A-C**). 4T1 tumors showed no significant changes in vessel density over time or after radiation. Irradiated 67NR tumors exhibited a decrease in vessel density compared to control tumors as soon as 1 day post-RT, which persisted after 10 days, at which point the vessel density was nearly half of the control tumors (**Figure 2A**). Despite the decrease in density, the cross sectional vessel area in irradiated 67NR tumors increased from 1 to 10 days, suggesting radiation-induced vascular remodeling over this timescale (**Figure 2D**). The average vessel size in 4T1 tumors was higher than 67NR vessels across all conditions but showed no significant differences over time or after irradiation. These observations correlated with US measurements of vascular index post-RT (**Figure 2E**). The correlation between IF and US-derived vascular measurements was lower at the 8/10 days post-RT timepoints likely due to the US imaging and IF sample collection occurring on different days. Despite this discrepancy, we observed a positive correlation between IF and US measurements, underscoring the feasibility of using US to identify and monitor vascular changes non-invasively over time. To further understand vascular effects and remodeling post-RT, we analyzed the changes in pericyte coverage of the vessels by measuring the fraction of CD31^+^ staining that also had ⍺SMA^+^ staining. Vessels in 67NR tumors exhibited an increase in pericyte coverage at 1 day post-RT that continued to increase up to 10 days post-RT (**Figure 2F-G**). In conjunction with the decline in vascularity and increase in vessel size, this demonstrates that vascular remodeling occurs in the 67NR tumors after a non-curative dose of RT. We did not observe any radiation-induced changes in pericyte coverage in 4T1 tumors. However, there was a decrease in coverage in both control and irradiated tumors from 1 to 10 days post-RT, likely due to the aberrant, disorganized vessel growth during tumor progression.

**Figure 2.**
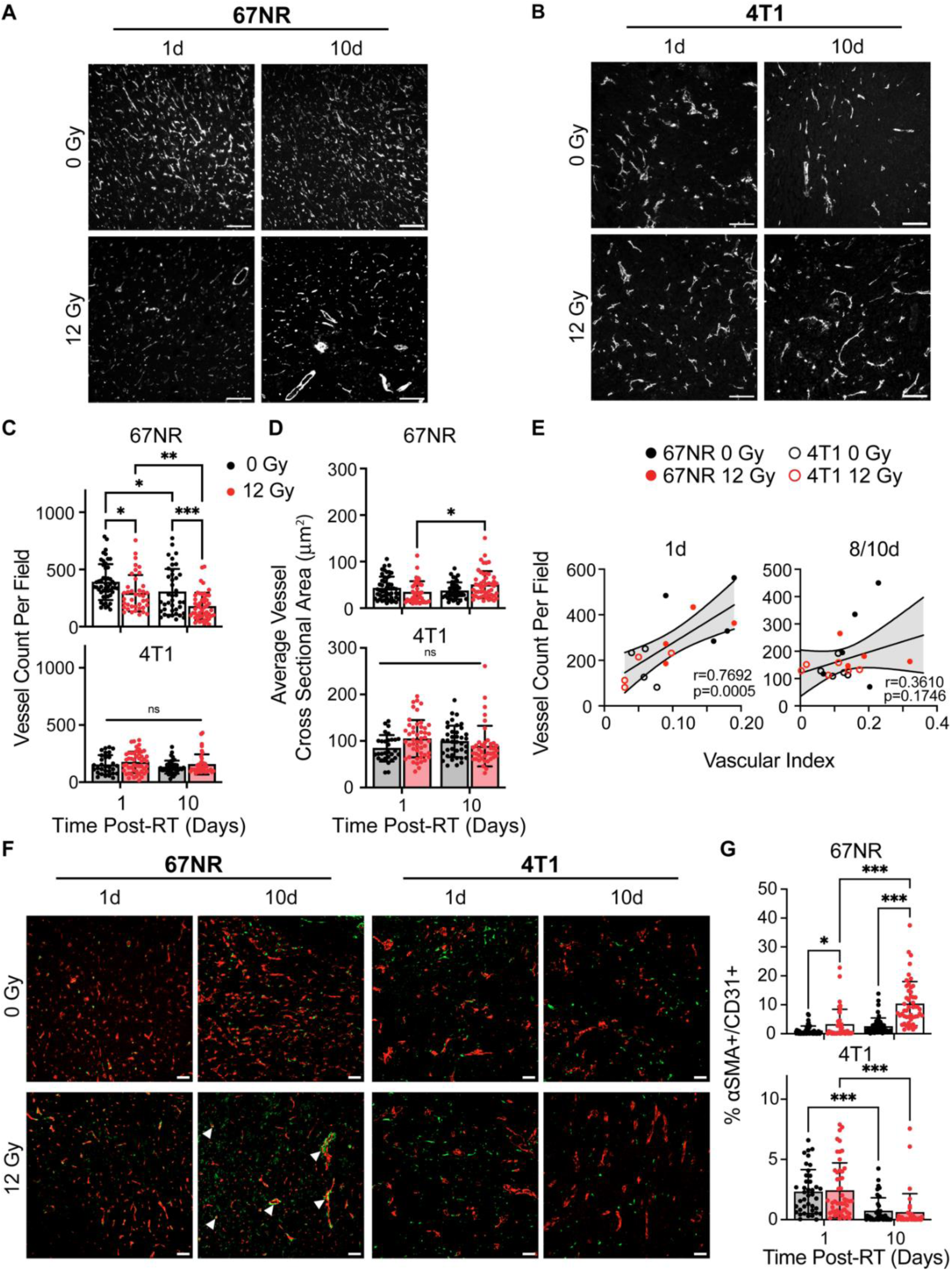
Immunofluorescence (IF) identifies vascular changes after radiation and between tumor types that correspond to US measurements. Representative CD31 IF staining of control and irradiated **(A)** 67NR and **(B)** 4T1 tumors at 1 and 10 days post-RT. **C.** Quantification of the number of CD31^+^ vessels per field of view (vessel density) in 67NR and 4T1 tumors. **D.** Quantification of the average cross-sectional vessel area per field of view in 67NR and 4T1 tumors. **E.** Pearson correlation analysis of vessel density (IF) and vascular index (US) quantification in 67NR and 4T1 tumors at 1 day and 8/10 days post-RT. Vascular index was measured at 8 days post-RT and compared to IF-derived vessel density in the same mouse at 10 days post-RT. Each point represents one mouse. **F.** Representative images of ⍺SMA and CD31 IF staining of control and irradiated tumors at 1 and 10 days post-RT. Arrows indicate pericyte-covered vessels. **G.** Quantification of the percent of CD31^+^ area that also contained ⍺SMA^+^ staining to identify pericyte-covered vessels. For each analysis, the number of biological replicates (n) were as follows: 67NR (0 Gy: n=5 mice for 1 day, n=4 mice for 10 day; 12 Gy: n=4 mice for 1 day, n=5 mice for 10 day 12 Gy), 4T1 (n=4 mice per condition and timepoint). Each data point represents one field of view with 10 fields of view per mouse. Scale bars are 100 μm. Error bars represent standard deviation with *p<0.05, **p<0.01, and ***p<0.001 as determined by two-way ANOVA.

### RT elicits a local increase in CD8^+^:CD4^+^ T cell ratios that follows vascular changes

We next sought to determine whether our non-curative dose of RT produced an immune response that correlated with the vascular changes observed during US imaging. Flow cytometry analysis of tumor-infiltrating myeloid and lymphocyte populations was performed at 1 and 10 days post-RT to identify early and intermediate responses in the local microenvironment. In both 67NR and 4T1 tumors, we did not see any significant changes in immune cells at 1 day post-RT. However, both tumor types showed significantly higher infiltration of CD45^+^ immune cells at 10 days post-RT (**Figure 3A**). Both irradiated and unirradiated 4T1 tumors showed an increase in immune cells—likely due to tumor growth and increased inflammation—while 67NR tumors showed a significant increase only in irradiated tumors, demonstrating the immune-stimulating effect from a single 12 Gy dose. While tumors showed no changes in the percentage of CD11b^+^ myeloid cells over time or between control and irradiated groups, the immune cells in 4T1 tumors were predominantly of myeloid lineage while 67NR tumors were comprised of approximately 50% myeloid cells (**Figure 3B**). No changes in macrophages, dendritic cells, and neutrophils in either tumor type over time or post-RT were observed (**Supplemental Figure S1**). Both 67NR and 4T1 tumors showed an increase in myeloid-derived suppressor cells between 1 and 10 days (**Supplemental Figure S1D, E**). However, this increase was abated by RT in 67NR tumors even after 10 days. 4T1 tumors also showed an increase in antigen presenting cells over time, though to a lesser extent in irradiated tumors.

**Figure 3.**
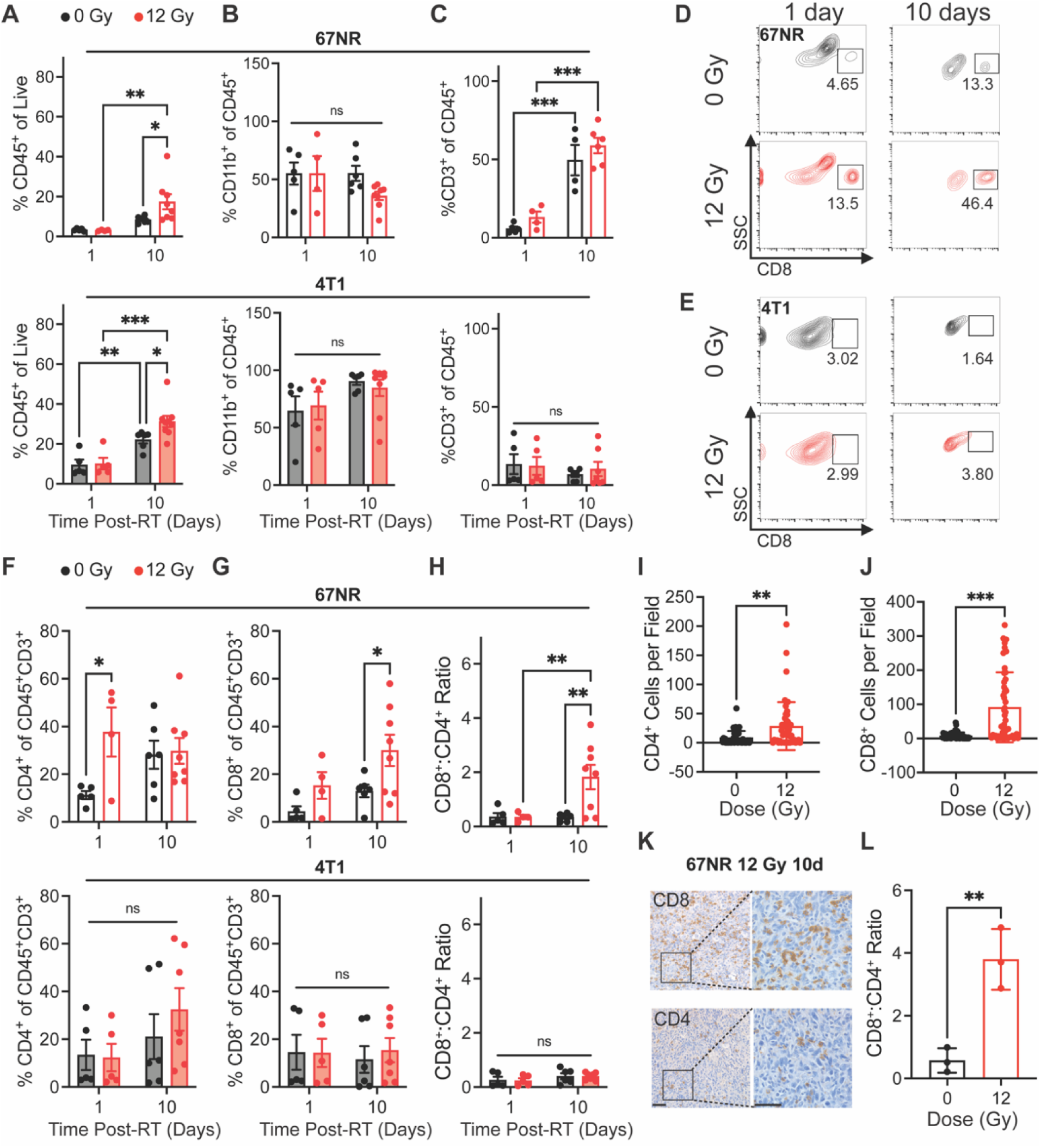
Immune cell infiltration post-RT differs between 67NR and 4T1 tumors. Flow cytometry characterization of tumor-infiltrating immune cell populations at 1 and 10 days post-RT, including **(A)** CD45^+^ immune cells, **(B)** CD11b^+^ myeloid cells, and **(C)** CD3^+^ T lymphocytes in 67NR (top; 0Gy: n=5 for 1 day, n=4 for 10 day; 12 Gy: n=4 for 1 day, n=6 for 10 day) and 4T1 (bottom; 0Gy: n=5 for 1 day, n=6 for 10 day; 12 Gy: n=5 for 1 day, n=7 for 10 day) tumors. Representative flow cytometry contour plots showing changes in CD8^+^ cytotoxic T cells after RT and over time in **(D)** 67NR and **(E)** 4T1 tumors. Further characterization of T lymphocyte subsets, including **(F)** CD4^+^ helper T cells, **(G)** CD8^+^ cytotoxic T cells, and **(H)** the ratio of CD8^+^:CD4^+^ T cells in 67NR (top) and 4T1 (bottom) tumors. Ratios calculated from absolute counts of CD45^+^CD3^+^CD4^+^ and CD45^+^CD3^+^CD8^+^ cells in each tumor. Quantification of IHC staining for **(I)** CD4^+^ helper and **(J)** CD8^+^ cytotoxic T cells in 67NR tumors. **K.** Representative images of CD4 and CD8 IHC staining in 12 Gy 67NR tumors at 10 days post-RT. Scale bars are 100 µm. **L.** Ratio of CD8^+^:CD4^+^ T cells calculated from IHC staining of 67NR tumors. Ratios calculated from the average number of cells per field in each condition. n=3 mice per condition. Error bars represent standard deviation with *p<0.05, **p<0.01, and ***p<0.001 as determined by two-way ANOVA.

An increase in CD3^+^ T cells in both irradiated and control 67NR tumors from 1 to 10 days post-RT suggests that the overall increase in immune cell populations after 10 days was driven by this population (**Figure 3C**). In 4T1 tumors, we observed no significant changes in total T cells over time or after radiation. Analysis of T cell subpopulations showed increased CD4^+^ T cell infiltration at 1 day post-RT in 67NR tumors that shifted to an increase in CD8^+^ T cell infiltration and CD8^+^:CD4^+^ T cell ratio at 10 days post-RT, which was not observed in 4T1 tumors (**Figure 3D-H**). This illustrates the immunostimulatory nature of our radiation scheme and differences in the expected immune landscapes between 67NR and 4T1 tumors [35–37]. We validated these results using IHC staining for CD8 in 67NR tumors, further confirming the ability of radiation to increase cytotoxic T cell infiltration 10 days post-RT (**Figure 3I-L**). Furthermore, the radiation-induced increase in the CD8^+^:CD4^+^ T cell ratio at 10 days post-RT showed an inverse relationship with the normalized vascular index (day 1/day -1) at 1 day post-RT (**Supplemental Figure S2**). Despite the variability observed across the irradiated mice, this inverse relationship suggests that a short-term reduction in vascular index and subsequent vascular normalization, which is characterized by increased pericyte coverage and improved drug perfusion and immune cell infiltration [38–40], may be linked to a later shift in the immune landscape of the tumor and highlights the potential utility of pairing non-invasive vascular monitoring with prediction of immune responses to RT.

### Systemic cytotoxic T cell response correlates with vascular changes and precedes local response to RT

Finally, we analyzed changes in immune cell populations in the spleen and lungs at 1 and 10 days post-RT to determine how systemic and distant responses to RT differ from the local response. Interestingly, 67NR-bearing mice exhibited a decrease in splenic CD4^+^ T cells and increase in CD8^+^ T cells 1 day post-RT, displaying a significant increase in the CD8^+^:CD4^+^ ratio at this timepoint (**Figure 4A-D**). These splenic cell populations returned to control proportions by 10 days post-RT, suggesting that the observed local shift in T cell populations in tumors at 10 days can be detected systemically and in the vascular index 24h after treatment (**Figure 1G**). In mice with 4T1 tumors, though we observed significant increases in splenic CD4^+^ T cells from 1 to 10 days post-RT, there was a decrease in splenic CD8^+^ T cells at 1 day post-RT and overall at 10 days post-RT, leading to a lower CD8^+^:CD4^+^ ratio at both timepoints in irradiated compared to control tumors (**Figure 4A-E**). IHC staining of spleens from mice with 67NR and 4T1 tumors at 10 days post-RT further corroborates our findings, showing negligible differences in CD4^+^ and CD8^+^ T cell infiltration between spleens from irradiated and control 67NR-bearing mice but a visible increase in the CD4^+^ T cell population in spleens from 4T1-bearing mice by 10 days post-RT (**Figure 4F-H**; **Supplemental Figure S3**). Analysis of myeloid cell subpopulations did not show significant differences in spleens from either 67NR-or 4T1-bearing mice after RT, though we observed increased proportions of macrophages, dendritic cells, and neutrophils from 1 to 10 days in 4T1-bearing mice (**Supplemental Figure S4**), likely due to splenomegaly.

**Figure 4.**
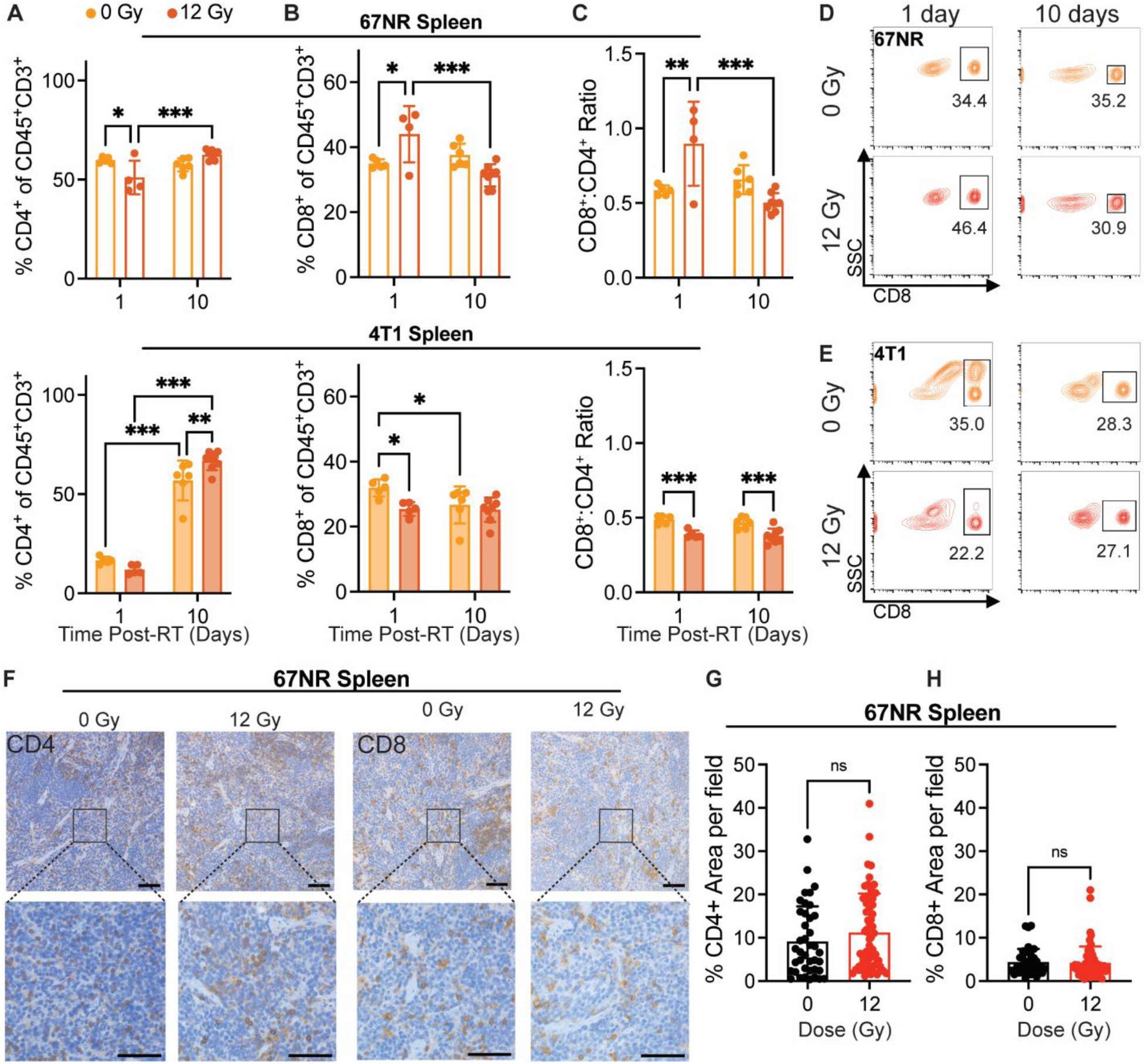
Splenic immune cell changes precede local tumor response to RT in 67NR tumors. Flow cytometry characterization of splenic immune cell populations at 1 and 10 days post-RT, including **(A)** CD4^+^ helper T cells, **(B)** CD8^+^ cytotoxic T cells, and **(C)** the ratio of CD8^+^:CD4^+^ T cells in spleens from mice with 67NR (top; 0 Gy: n=5 for 1 day, n=4 for 10 day; 12 Gy: n=4 for 1 day, n=6 for 10 day) and 4T1 (bottom; 0 Gy: n=5 for 1 day, n=6 for 10 day; 12 Gy: n=5 for 1 day, n=7 for 10 day) tumors. Ratios calculated from absolute counts of CD45^+^CD3^+^CD4^+^ and CD45^+^CD3^+^CD8^+^ cells in each spleen. Representative contour plots showing changes in splenic CD8^+^ T cells after radiation and over time in mice with **(D)** 67NR and **(E)** 4T1 tumors. **F.** Representative images of CD4^+^ and CD8^+^ IHC staining in the spleens of 67NR-bearing mice at 10 days post-RT. Scale bars are 100 µm. Quantification of the percent area of **(G)** CD4^+^ and **(H)** CD8^+^ staining per field of view in spleens from 67NR-bearing mice at 10 days post-RT. n= 3 mice per condition. Each point represents one field of view with 10 total fields per mouse. Error bars represent standard deviation with *p<0.05, **p<0.01, and ***p<0.001 as determined by two-way ANOVA.

Quantification of immune cells in the lungs showed no changes after radiation for either tumor model. However, clear differences in the immune cell populations between lungs in mice with 67NR and 4T1 tumors were detected. While the lungs of mice with 67NR tumors showed no significant increases in CD45^+^ immune cells over time, the lungs of 4T1-bearing mice exhibited significant increases under both control and irradiated conditions (**Supplemental Figure S5A-D**). The immune subpopulations present in the lungs mirror those in the tumor microenvironment, with lungs dominated by CD11b^+^ myeloid cells in mice with 4T1 tumors and a substantially higher proportion of CD3^+^ T cells in mice with 67NR tumors (**Supplemental Figure S5E-I**). Analysis of myeloid subpopulations in the lungs also show overall higher proportions in mice with 4T1 compared to 67NR tumors with the largest differences occurring over time rather than after radiation (**Supplemental Figure S5E**). Since 4T1 cells readily metastasize to the lungs but 67NR cells are not metastatic [41], these differences in the immune landscape are expected. Within each tumor type, the consistency of the immune landscape between control and irradiated lungs highlight that our radiation scheme does not impact distant sites. Taken together, our non-curative dose of RT induces retraction in the vasculature of immunostimulatory hot tumors (67NR) that corresponds to increased vessel maturity and CD8^+^:CD4^+^ T cell ratios while immunosuppressive cold tumors (4T1) show no change in vascular structure and thus no change in the CD8^+^ T cell landscape (**Figure 5**). This local change in immune microenvironment at 10 days post-RT is seen systemically as early as 1 day post-RT, where splenic CD8^+^:CD4^+^ T cell ratios are elevated in mice bearing 67NR tumors.

**Figure 5.**
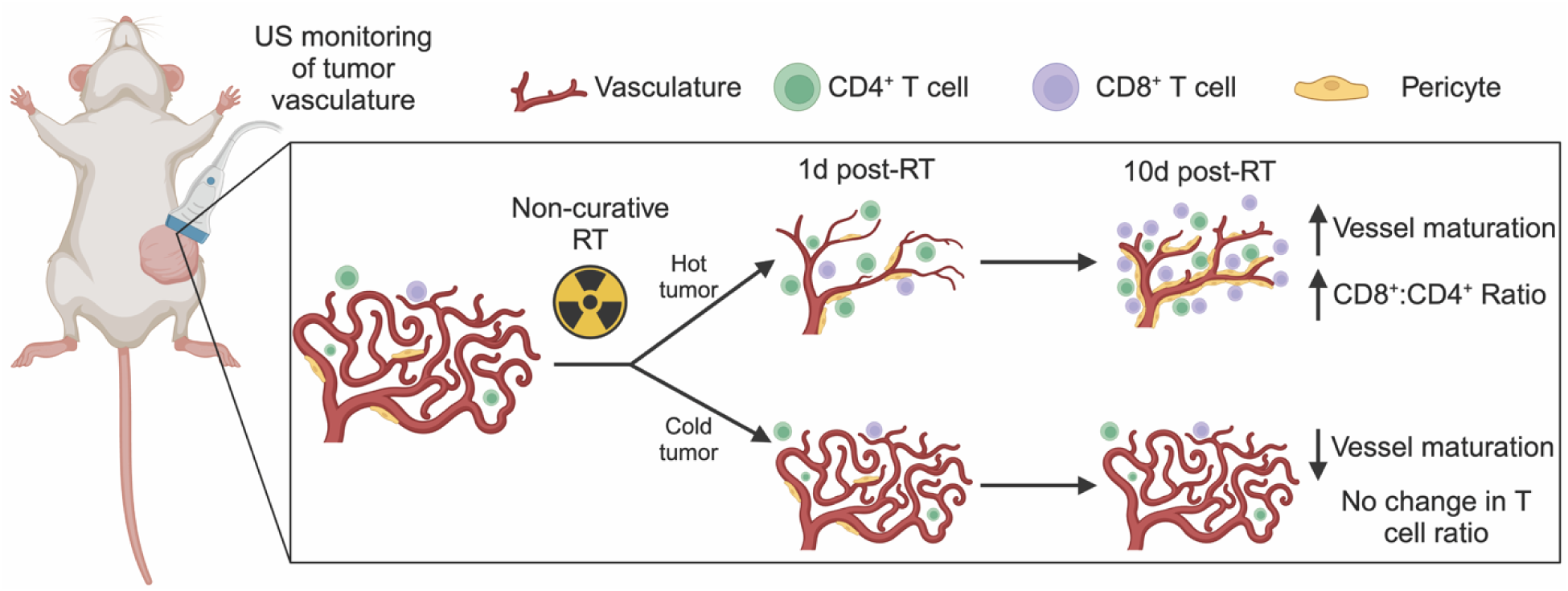
US detection of vascular changes post-RT corresponds to shifts in the immune microenvironment. Ultrafast power doppler US reveals that non-curative RT induces retraction and then maturation of vasculature that is linked to an increase in the number and proportion of CD8^+^ T cells in immunostimulatory hot tumors. In contrast, immunosuppressive cold tumors experience no gross changes in tumor vasculature post-RT but exhibit a decrease in pericyte coverage that corresponds to no change in CD8^+^ T cells within the tumor.

## Discussion

With recent progress in the field of immunotherapy, the limited treatment options for TNBC patients may be expanded through personalized medicine approaches that combine prognostic monitoring with new treatments. Non-invasive methods of predicting and tracking tumor response to therapy have the potential to greatly improve patient outcomes. The tumor vasculature is particularly important in this regard as it can give insight into the degree of malignancy and predict drug delivery efficacy and immune cell infiltration to the tumor. In this proof-of-principle study, we used ultrafast power doppler US imaging to track RT-induced vascular changes over time in two breast cancer models that differ in metastatic capacity and immunogenicity [35–37]. First, we showed that ultrafast power doppler US can non-invasively monitor post-RT vascular changes that correspond to vessel characteristics as determined by IF (**Figure 2**). We then identified that the decrease in vascular index observed via US at 1 day post-RT in 67NR mice corresponded to a splenic increase in CD8^+^:CD4^+^ ratio at 1 day post-RT and preceded a significant local increase in the CD8^+^ cytotoxic T cell population that was only observed in irradiated tumors (**Figure 3**). Together, these data establish our US technique as an accurate method to track tumor vasculature changes over time and a potential proxy for predicting both systemic and local immune response to RT.

Prior work has tracked the tumor vascular response to radio-and immunotherapy using high-frequency (>15MHz) power doppler US without contrast, such as El Kaffas et al. who identified the same trend of decreased vasculature 1 day after irradiating MDA-MB-231 human TNBC tumors with 8 and 16 Gy [26]. More recently, similar reductions in vasculature have also been observed using contrast enhanced super harmonic imaging methods that transmit with low frequency pulses and receive at super harmonic frequencies, which allows for high resolution visualization of vasculature [23]. US localization microscopy is an emerging high frame rate contrast-enhanced technique for tracking the trajectories of individual microbubbles [42]. Opacic et al. demonstrated that super resolution imaging is capable of quantifying tumor at the individual vessel level but requires longer acquisition time [43]. Our study combined contrast enhancement provided by microbubbles and high acquisition rate of planewave ultrafast doppler US without employing localization methods. Our method operated at clinical frequency (<15MHz), was implemented with a commercial linear array, and only required a small bolus injection of microbubbles. This system achieved *in vivo* imaging with sub-millimeter level resolution laterally and axially sufficient for visualizing and quantifying rodent microvessels over 15 mm below the tissue surface [27]. Out-of-plane sampling of the tumor could improve 3D imaging by decreasing the spacing to a step size lower than one half the elevational resolution of the transducer [44] or using rotational compounding could potentially be performed as in Demené et al. [28]. Additionally, while our US measurements of vascular index correlated strongly with IF vessel density at 1 day post-RT when timepoints for each assay were matched, we observed a weaker correlation between our later timepoints where US and IF data were not collected on the same day due to technical challenges with sample processing. Future studies will be needed to optimize processing time in order to collect US, IF, and flow cytometry data on the same day. Still, microvascular imaging with US is sensitive to changes that occur during the response to cancer therapy and could help inform guidance criteria, such as RECIST (Response Evaluation Criteria in Solid Tumors) [45], by providing non-invasive measurements of functional activity that correspond to known biological responses to radio-or immunotherapy.

Previous work has found the vasculature of 67NR tumors to be substantially different from 4T1 tumors [30,46], which was corroborated by our study. The slower growing 67NR tumors form vessels with a thicker endothelium compared to the 4T1 tumors, which are characterized by leaky, tortuous vasculature and significant areas of necrosis. RT is known to cause both acute, direct damage to tumors in the form of DNA double-strand breaks in addition to indirect damage through the generation of reactive oxygen species and recruitment of immune cells that can occur days to weeks following exposure [47,48]. Based on our US and IHC data, it is likely that the direct damage caused by RT within minutes after exposure caused a retraction of the mostly intact 67NR vasculature by 24 hours post-RT—an effect seen after single doses of RT [49,50]—while the less vascularized 4T1 tumors were largely unaffected by our RT scheme. Because 4T1 tumors are necrotic, immunologically cold, and highly aggressive relative to 67NR tumors [36,37], a higher dose or fractionated regimen of RT may be necessary to elicit vascular changes and impact immune cell infiltration. While the overall vessel density remained low in the 67NR tumors through 10 days post-RT, we observed a shift toward larger vessels and an increase in pericyte coverage throughout the irradiated tumors that were not seen in the control. Additional US analyses to quantify the number of larger vessels and further characterize the physical aspects of the vascular network are required to support these findings. However, this initial observation indicates radiation-induced vascular remodeling in agreement with previous studies that show reduced microvascular density, larger vessel size, and increased pericyte coverage within days after exposure to doses at or above 12 Gy [21,51,52].

Traditionally high, fractionated doses of RT have been reported to cause vascular retraction and hypoxia that recruits myeloid cells and ultimately lead to vascular regrowth and tumor recurrence [48,53], highlighting the importance of dosing in both vascular and immune cell changes within tumors. While further studies are needed to determine the optimal dose and fractionation schemes that elicit vascular changes and lead to favorable immune cell infiltration, the non-ablative RT scheme employed in our study induces vascular normalization that has been shown in pancreatic cancer leading to recruitment and infiltration of T lymphocytes that improve tumor immunogenicity [39,40]. We observed a significant increase in the CD8^+^:CD4^+^ T cell ratio only in the irradiated 67NR tumors, which also exhibited a reduction in vessel density and increase in vessel size and pericyte coverage. The mature vessels detected in the irradiated 67NR tumors may also contribute to the increase in CD8^+^ T cell infiltration via upregulation of adhesion molecules, which are often downregulated on tumor endothelial cells that line the vasculature and therefore prevent transendothelial migration of lymphocytes from circulation into the tumor [54,55]. Additional US and histological analyses are needed to confirm the direct association between immune cell infiltration and vascular changes in our model. Future studies will characterize the adhesion molecule expression and permeability of the vasculature following RT as well as include vascular disrupting agents (e.g. combretastatin A4 phosphate [56]) to identify the direct effects of vascular damage on our observed shifts in the immune microenvironment.

We also observed a significant increase in the splenic CD8^+^:CD4^+^ T cell ratio of mice with 67NR tumors at 1 day post-RT. This observation is indicative of a systemic response elicited by local radiation, thus agreeing with previous reports of this phenomena in pre-clinical studies of RT and immunotherapy [57,58]. While the exact link between changes in splenic and tumor immune cell populations has not been defined, the spleen is a critical site of immune tolerance to induce anti-tumor immunity [59,60]. Our findings in the splenic response to local tumor irradiation therefore illustrate the potential of RT to be combined with immunotherapy and the need for a method to track and predict the success of combination treatments. The observed vascular change in response to RT in 67NR tumors using non-invasive US imaging could be used to determine immune cell infiltration and response to therapy as we observed substantial vascular changes after 1 day but no immune cell changes until 10 days post-RT. A recent study correlated low vasculature and high inflammation in basal cell carcinomas with improved therapeutic outcomes [24], which underscores the feasibility of using the vascular response to predict immune infiltration. CT imaging biomarkers have also been developed to correlate tumor-immune microenvironment changes with response to immune checkpoint blockade [61], further demonstrating the utility of coupling non-invasive tumor monitoring with evaluating the immune response.

## Conclusion

Identifying methods that accurately monitor changes in tumor vasculature over time and non-invasively after therapy may be critical not only for assessing the response to adjuvant immunotherapy and determining outcomes early in the treatment process but also for guiding clinical treatment strategies. Future experiments will provide more in depth characterization of vascular changes using US and further link the tumor vasculature and changes in tumor-infiltrating immune cell populations. This work demonstrates the potential for using ultrafast US imaging as a personalized medicine technique to evaluate changes in the tumor microenvironment following RT. Monitoring vascular changes to predict therapy response and shifts in the immune landscape of the tumor may improve outcomes for patients with limited treatment options.

## Abbreviations

⍺SMA: ⍺-smooth muscle actin
DAPI: 4’,6-diamidino-2-phenylindole dihydrochloride
Gy: Gray
IQ: in-phase and quadrature
IF: Immunofluorescence
IHC: Immunohistochemistry
NBF: Neutral buffered formalin
PBS: phosphate-buffered saline
RECIST: Response Evaluation Criteria in Solid Tumor
RF: Radiofrequency
RT: radiotherapy
SVD: Single value decomposition
TNBC: Triple negative breast cancer
US: Ultrasound

## Acknowledgements

The authors thank Dr. Michael Freeman for irradiator use and Dr. Austin Kirschner and the radiation physics staff for constructing the Cerrobend shield. We also thank the Vanderbilt Translational Pathology Shared Resource (TPSR) supported by NCI/NIH Cancer Center Support Grant P30CA068485 for tissue sample preparation. This research was financially supported by the Focused Ultrasound Foundation Cancer Immunotherapy Award and NIH grant R00CA201304.

## Contributions

M.R. and C.F.C conceptualized the study and supervised the project. J.K., J.X., A.H., A.S., and C.F.C developed the US protocol. S.E.M., J.K., J.X., A.H., B.C.H., M.A.M., A.S., and E.J.L. performed experiments. S.E.M. and J.X. performed data analysis and wrote the manuscript. S.E.M., J.X., J.K., C.F.C, and M.R. revised the manuscript.

