## Supplementary Figures for "Ultrafast Power Doppler Ultrasound Enables Longitudinal Tracking of Vascular Changes that Correlate with Immune Response After Radiotherapy"

### Supplemental Figures

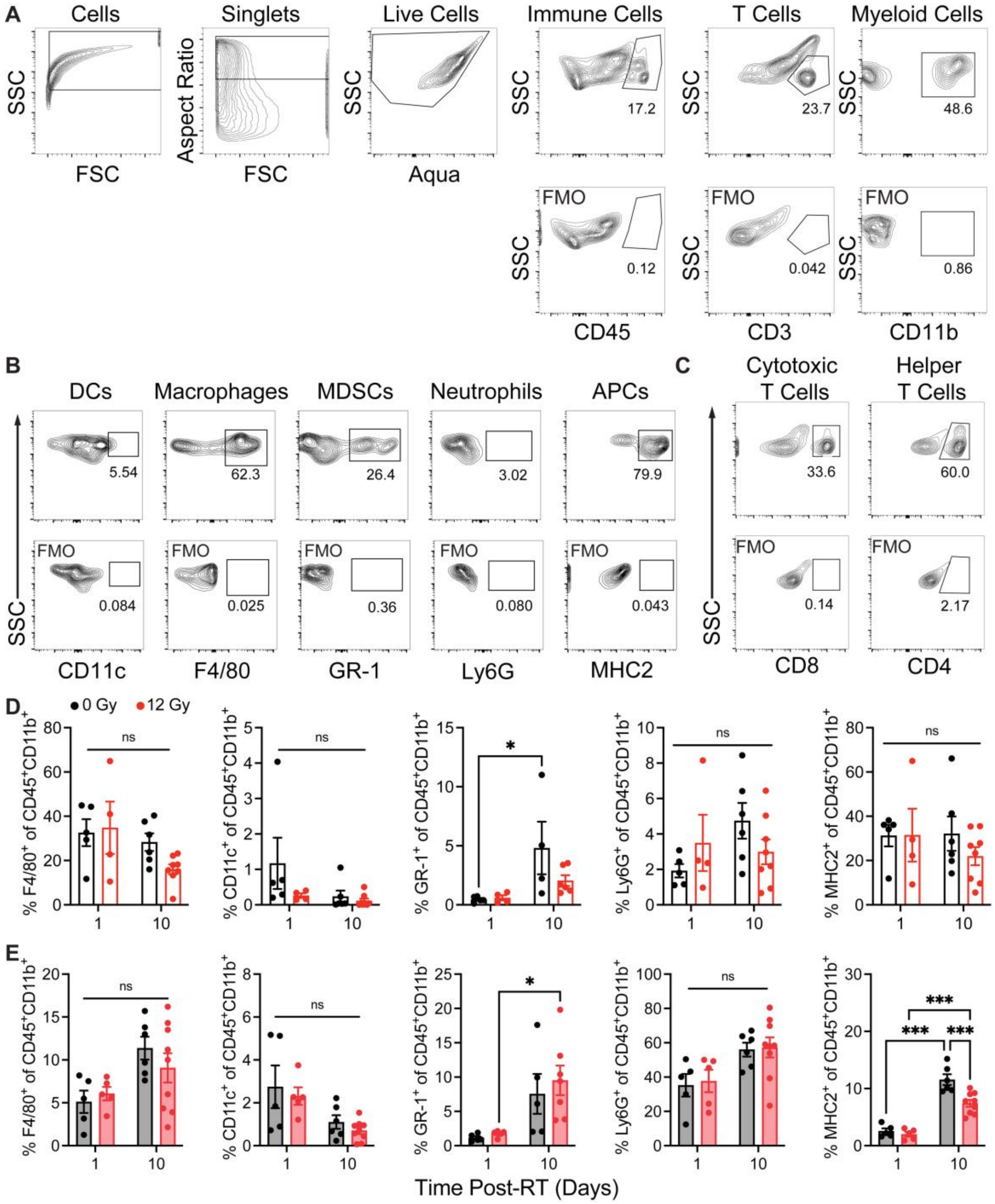


**Figure S1. Tumor-infiltrating immune cell flow cytometry analysis. A.** Full gating strategy for myeloid and T cells in 67NR and 4T1 tumors with full minus one (FMO) controls shown in the bottom panels. **B.** Representative gating for myeloid subpopulations. **C.** Representative gating for T cell subpopulations. Percentage of myeloid subpopulations, including F4/80$^+$ macrophages, CD11c$^+$ dendritic cells, GR-1$^+$ myeloid-derived suppressor cells, Ly6G$^+$ neutrophils, and MHC2$^+$ antigen presenting cells, in **(D)** 67NR tumors (0 Gy: n=5 for 1 day, n=4 for 10 day; 12 Gy: n=4 for 1 day, n=6 for 10 day) and **(E)** 4T1 tumors (0 Gy: n=5 for 1 day, n=6 for 10 day; 12 Gy: n=5 for 1 day, n=7 for 10 day) 1 and 10 days after 12 Gy RT exposure. Error bars represent standard deviation with *p<0.05 and ***p<0.001 as determined by ANOVA.

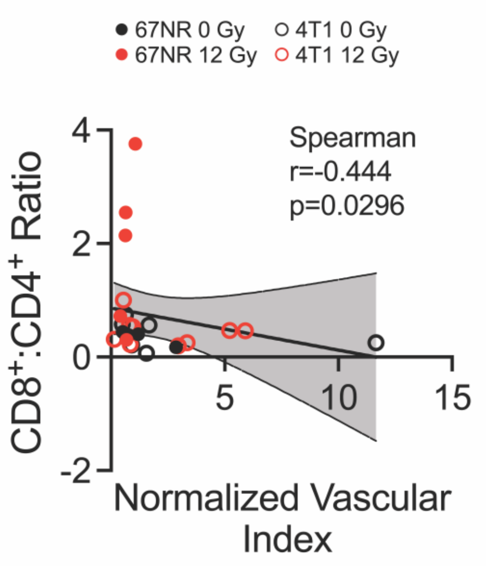


**Figure S2. CD8$^{+}$:CD4$^{+}$ T cell ratio 10 days post-RT inversely trends with vascular index 1 day after RT.** Spearman correlation and linear regression analyses of the 10 day post-RT tumor CD8$^{+}$:CD4$^{+}$ T cell ratio and 1 day post-RT vascular index (normalized to day -1). Each point represents one mouse with n=4 mice for 67NR 0 Gy, n=5 mice for 67NR 12 Gy, n=6 mice for 4T1 0 Gy, and n=9 mice for 4T1 12 Gy.

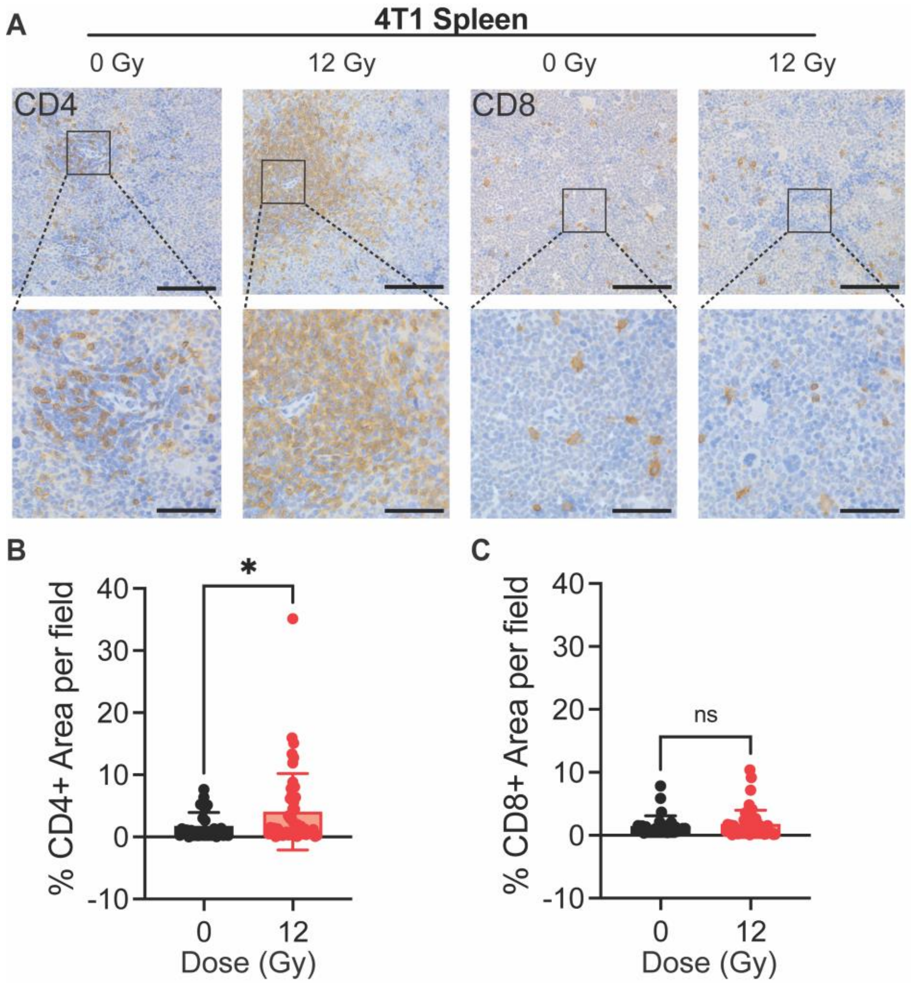


**Figure S3. Immunohistochemistry (IHC) validates T cell population shifts in spleens from mice with 4T1 tumors post-RT. A.** Representative images of $CD4^+$ and $CD8^+$ IHC staining of spleens in mice with 4T1 tumors at 10 days post-RT. Scale bars are 100µm (top row) and 50 µm (bottom row). Quantification of the percent area of **(B)** $CD4^+$ and **(C)** $CD8^+$ staining in spleens at 10 days post-RT. n=3 mice per condition. Each point represents one field of view with 10 total fields per mouse. Error bars indicate standard deviation with *p<0.05 determined by unpaired t-test.

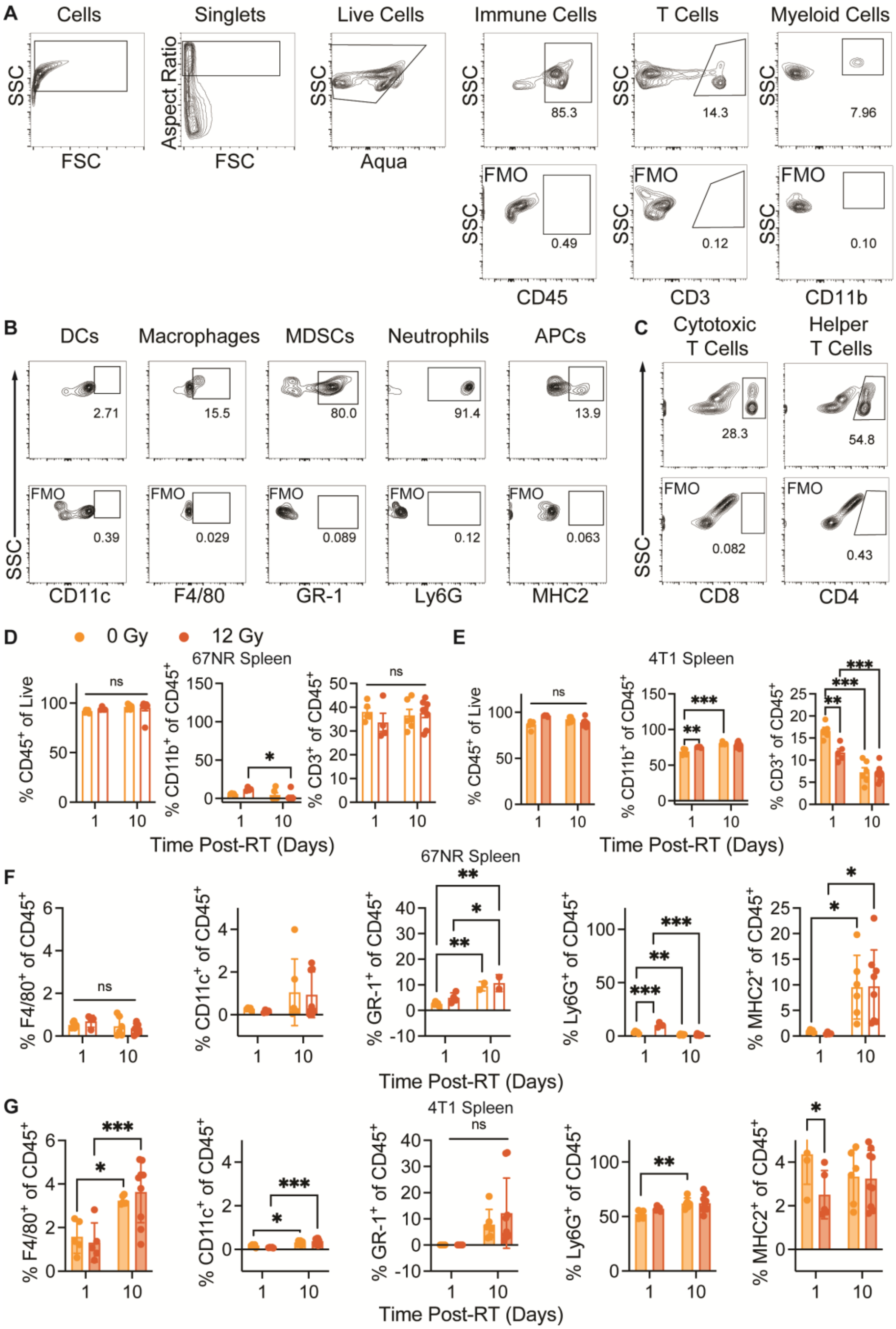


**Figure S4. Flow cytometry analysis of splenic immune cell populations. A.** Full gating strategy for myeloid and T cells in spleens with full minus one (FMO) controls shown in the bottom panels. **B.** Representative gating for myeloid subpopulations. **C.** Representative gating for T cell subpopulations. Percentage of CD45$^+$ immune, CD11b$^+$ myeloid, and CD3$^+$ T cell populations in spleens from **(D)** 67NR and **(E)** 4T1-bearing mice 1 and 10 days after 12 Gy RT exposure of the tumor. Percentage of myeloid subpopulations in the spleens of **(F)** 67NR and **(G)** 4T1-bearing mice, including F4/80$^+$ macrophages, CD11c$^+$ dendritic cells, GR-1$^+$ myeloid-derived suppressor cells, Ly6G$^+$ neutrophils, and MHC2$^+$ antigen presenting cells. For 67NR, 0 Gy: n=5 mice for 1 day, n=4 for 10 day; 12 Gy: n=4 for 1 day, n=6 for 10 day. For 4T1, 0 Gy: n=5 mice for 1 day, n=6 for 10 day; 12 Gy: n=5 for 1 day, n=7 for 10 day. Error bars represent standard deviation with *p<0.05, **p<0.01, and ***p<0.001 as determined by ANOVA.

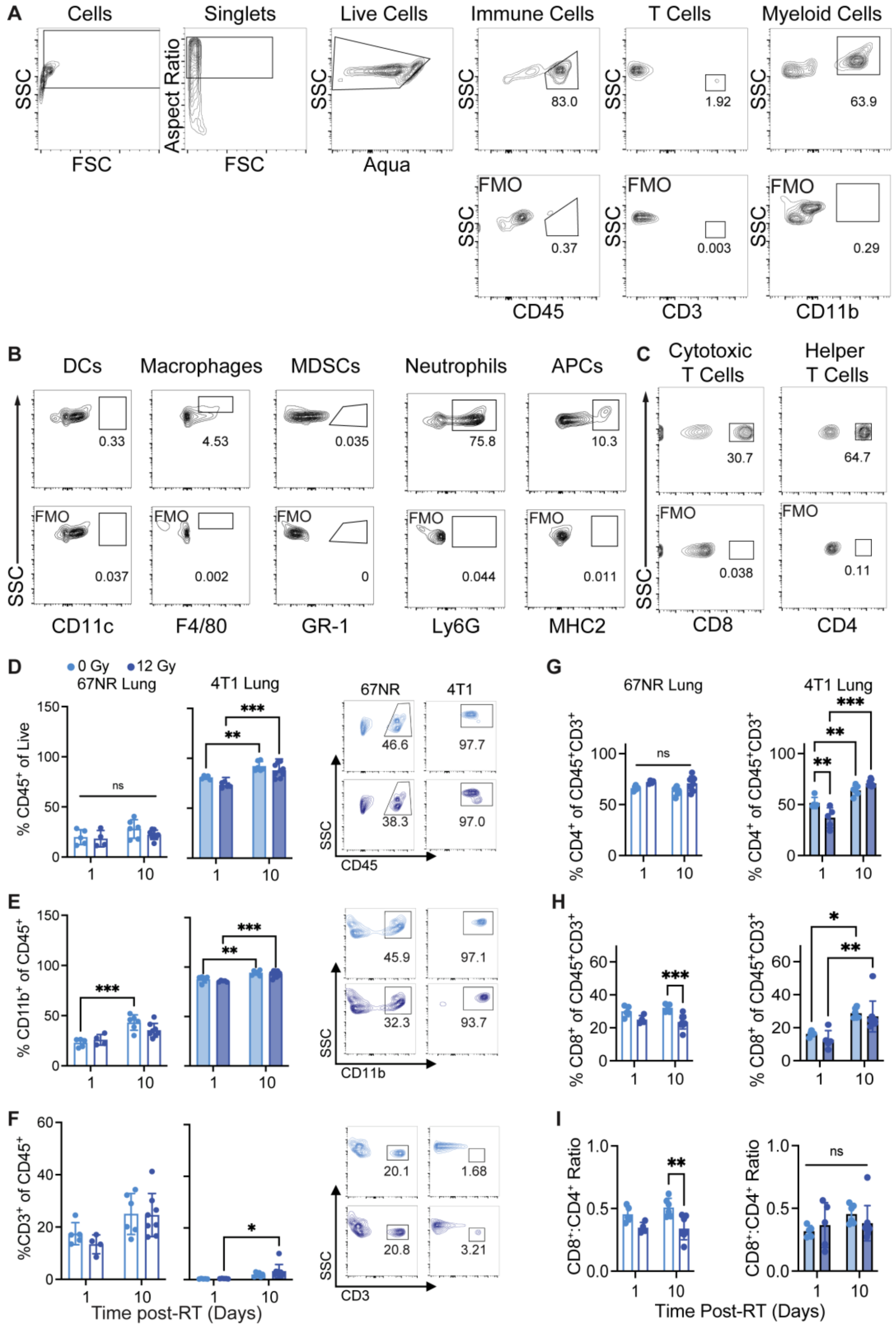


**Figure S5. Flow cytometry analysis of lung immune cell populations. A.** Full gating strategy for myeloid and T cells in lungs with FMO controls shown in the bottom panels. **B.** Gating strategy for myeloid subpopulations. **C.** Gating strategy for T cell subpopulations. Flow cytometry characterization and representative contour plots of lung immune cell populations at 1 and 10 days post-RT, including **(D)** $CD45^+$ immune cells, **(E)** $CD11b^+$ myeloid cells, and **(F)** $CD3^+$ T cells in lungs of mice with 67NR (left; 0Gy: n=5 for 1 day, n=4 for 10 day; 12 Gy: n=4 for 1 day, n=6 for 10 day) and 4T1 (right; 0 Gy: n=5 for 1 day, n=6 for 10 day; 12 Gy: n=5 for 1 day, n=7 for 10 day) tumors. Error bars represent standard deviation with *$p<0.05$, **$p<0.01$, and ***$p<0.001$ as determined by two-way ANOVA. Percentage of **(G)** $CD4^+$ helper T cells, **(H)** percentage of $CD8^+$ cytotoxic T cells, and **(I)** ratio of $CD8^+$:$CD4^+$ T cells calculated from absolute counts of $CD45^+CD3^+CD4^+$ and $CD45^+CD3^+CD8^+$ cells in the lungs of mice bearing 67NR and 4T1 tumors lungs 1 and 10 days after 12 Gy RT exposure of the tumor. For 67NR, 0 Gy: n=5 mice for 1 day, n=4 for 10 day; 12 Gy: n=4 for 1 day, n=6 for 10 day. For 4T1, 0 Gy: n=5 mice for 1 day, n=6 for 10 day; 12 Gy: n=5 for 1 day, n=7 for 10 day. Error bars represent standard deviation with *$p<0.05$, **$p<0.01$, and ***$p<0.001$ as determined by ANOVA.

**Table S1. Linear Regression Analysis[a].**

| Condition | Slope | | Intercept | | n |
|---|---|---|---|---|---|
| | m | 95% CI | m | 95% CI | |
| 67NR 0 Gy | 0.739 | 0.528 to 0.950 | 17.11 | -41.46 to 75.68 | 22 |
| 67NR 12 Gy | 0.237 | 0.139 to 0.334 | 92.84 | 68.23 to 117.4 | 20 |
| 4T1 0 Gy | 0.865 | 0.608 to 1.122 | -28.94 | -95.31 to 37.43 | 32 |
| 4T1 12 Gy | 0.716 | 0.501 to 0.932 | -4.959 | -66.44 to 56.53 | 39 |

[a]m=mean; CI=confidence interval; n=number of measurements with 9 mice included in each of the 67NR 0 and 12 Gy groups, 15 in 4T1 0 Gy, and 13 in 4T1 12 Gy
